# Expression of LTR and LINE1 transposable elements defines atypical teratoid/rhabdoid tumor subtypes

**DOI:** 10.1101/2025.05.13.653713

**Authors:** Martin V. Hamann, Shweta Godbole, Maisha Adiba, Sabrina M. Leddy, Jelena Navolić, Ghazaleh Tabatabai, Daniel J. Merk, Ulrike C. Lange, Julia E. Neumann

**Affiliations:** Leibniz Institute of Virology (LIV), Hamburg, Germany; Center for Molecular Neurobiology (ZMNH), University Medical Center Hamburg–Eppendorf, Hamburg, Germany; Department of Molecular Biology and Genetics, Cornell University, Ithaca, NY, USA; Department of Neurology and Interdisciplinary Neuro-Oncology, Hertie Institute for Clinical Brain Research, University Hospital Tübingen, Eberhard Karls University, Tübingen, Germany; Institute for Infection Research and Vaccine Development, University Medical Center Hamburg–Eppendorf, Hamburg, Germany; Institute of Neuropathology, University Medical Center Hamburg–Eppendorf, Hamburg, Germany

**Keywords:** transposable elements, atypical teratoid/rhabdoid tumors (ATRTs), DNA methylation data, LINE1, LTR elements, transcriptome

## Abstract

Atypical teratoid rhabdoid tumors (ATRTs) are aggressive central nervous system tumors mainly affecting young children. Extensive molecular characterization based on gene expression and DNA methylation patterns has solidly established three major ATRT subtypes (MYC, SHH and TYR), which show distinct clinical features, setting the basis for more effective, targeted treatment regimens. Transcriptional activity of transposable elements (TEs), like LINE1s and LTRs, is tightly linked with human cancers as a direct consequence of lifting epigenetic repression over TEs. The sole recurrent biallelic loss-of-function mutation in *SMARCB1* in ATRTs, a core component of the SWI/SNF chromatin remodeling complex, raises the question of how TE transcription contributes to ATRT development.

Here, we comprehensively investigate the transcriptional profiles of 1.9M LINE1 and LTR elements across ATRT subtypes in primary human samples. We find TE transcription profiles are unique, allowing sample stratification into ATRT subtypes. The TE activity signature in ATRT-MYC subtype is unique, setting these tumors apart from SHH and TYR ATRTs. More specifically, ATRT-MYC shows broadly reduced transcript levels of LINE1 and ERVL-MaLR subfamilies. ATRT-MYC is also unique in having significantly less LTR and LINE1 loci with bidirectional promoter activity. Furthermore, we identify 849 differentially transcribed TEs in primary samples, which are predictive towards established ATRT-SHH and-MYC cell line models.

In summary, including TE transcription profiles into the molecular characterization of ATRTs might reveal new tumor vulnerabilities leading to novel therapeutic interventions, such as immunotherapy.

## Material and Methods

### RNA Sequencing

Transcriptome datasets for ATRT cell lines BT12, BT16, CHAL.02 and CHLA.06 were generated as described previously [1]. In brief, total RNA was isolated from cell pellets and sample quality was assessed with the Agilent 2100 Fragment Analyzer total RNA kit (Agilent Technologies, Inc., US). Samples with RIN >9 were selected for library construction. Libraries were prepared using the NEBNext Ultra II Directional RNA Library Prep kit with 200ng of total RNA input and polyA selection according to the manufacturer’s instructions. All libraries were sequenced on the Illumina NovaSeq 6000 platform in paired-end mode with 2×50bp reads and at a depth of approx. 25 million clusters each. Data is accessible via GEO platform accession GSE231287.

### Public datasets and genome references

Raw transcriptome, DNA methylome and meta data for human primary ATRT samples were compiled from Andrianteranagna et al. and Lobón-Iglesias et al. [2], [3] using SRA database accession numbers SRP457328 & SRP322114. We re-analyzed raw transcriptome and DNA methylation data against the human genome reference build GRCh38 (hg38) as described below. TE information was obtained from the UCSC genome browser RepeatMasker track (Assembly: Human Dec. 2013 (GRCh38/hg38); Data last updated at UCSC: 2022-10-18) and filtered for class LINE1 and LTR elements [4]. A summary of element counts and family subtyping are provided in Supplementary Tables 1 and 2.

### DNA methylome analysis

Idat files were processed in R (Version 4.4.2). The files were read using the minfi package (Version 1.51.1) [5]. Differentially and most variably methylated probes/CpG sites were found using the limma package (Version 3.61.2) [6], correcting multiple testing using Benjamini Hochberg (cut-off of 5% FDR). Beta-values of the 20,000 most variable CpG sites were selected for further analysis. Heatmaps were made using the ComplexHeatmap package (Version 2.22.0) [7].

### Gene expression analysis

Gene expression analysis is based on read mapping utilizing Salmon with standard settings (--validateMappings) (v1.10.1) [8]. Downstream analysis was conducted with packages DESeq2 (v1.44.0) [9], pheatmap (v1.0.12) [10], ComplexHeatmap (v2.20.0) [11], ggplot2 (v3.5.1) [12], EnhancedVolcano (v1.22.0) [13], org.Hs.eg.db (v3.19.1) [14] and limma (v3.60.3) [6] in R (v4.4.0).

### TE transcriptional profiling

Transcription profiles of LINE1 and LTR elements were obtained with the SalmonTE tool using standard parameter--validateMappings and--minScoreFraction 0.65 [8], [15]. Downstream analysis was conducted as above with packages DESeq2 (v1.44.0) [9], pheatmap (v1.0.12) [10], ComplexHeatmap (v2.20.0) [11], ggplot2 (v3.5.1) [12], EnhancedVolcano (v1.22.0) [13], org.Hs.eg.db (v3.19.1) [14] and limma (v3.60.3)) [6] in R (v4.4.0). To assess the TE transcription at subfamily level, relative normalized TE counts were summarized retaining only the elements with at least 5 mapped reads on average in at least one ATRT subtype. The expression score was calculated based on the average expression of each subfamily using matrixStats (v1.5.0), and the occurrence score was defined as the number of expressed loci per subfamily.

### Bidirectional TE transcription analysis

*Quantification of sense and antisense transcription:* To quantify sense and antisense transcription at individual TE loci, the pre-trimmed, paired-end RNAseq reads were mapped with SQuIRE to the hg38 assembly, which uses STAR (2.5.3a) as an aligner [16], [17]. The resulting BAM files containing strand-assigned reads were processed using featureCounts (subread, v2.0.3) to count sense and antisense reads [18]. A GTF file of the hg38 RepeatMasker annotation was used to provide the locus-level coordinate and strand information to featureCounts. We used the-s option to engage strand filtration, meaning that featureCounts compared the reads mapped to each TE locus to the GTF annotation, counting both sense and antisense reads per loci. The -- fraction &-M options in featureCounts were used to account for multi-mapping, meaning that each reported alignment from a multi-mapping read carries a fractional count of 1/x, instead of 1, where x is the total number of alignments reported for the same read. We then subset these raw read counts, removing simple repeats, SINE elements, and DNA transposons. We also removed TE loci within gene exons. In the final array, we retained intronic and intergenic LINE1 and LTR loci. This raw counts table was normalized using DESeq2 (v1.40.2) [9].

*Detection of bidirectional transcription:* To identify bidirectionally transcribed LTR and LINE1 loci within each group (MYC, SHH, TYR), we filtered for loci with ≥10 DESeq2-normalized reads in both the sense and antisense orientations. To minimize false positives, this threshold had to be met in at least 50% of the samples within each group. Loci with <10 reads in both orientations were discarded. Loci with <10 reads in only one orientation were classified as either sense-transcribed (>10 sense reads, <10 antisense reads) or antisense-transcribed (>10 antisense reads, <10 sense reads), provided it met the 50% sample threshold. For further statistical analyses and visualization, we extracted LINE1 and LTR loci separately, labelled by group as sense, antisense, and bidirectionally transcribed categories. When comparing groups, loci which were sorted into different categories between groups were labelled as ‘mixed’. ANOVA tests were performed in R (v4.3.1). For visualization, box plots were created in R using ggplot2 (v3.4.2).

## Introduction

Atypical teratoid rhabdoid tumors (ATRTs) are aggressive pediatric malignancies that manifest in the central nervous system (CNS) [19]. They belong to the rhabdoid tumor (RT) family, which is characterized by biallelic inactivation of *SMARCB1* in a genome with otherwise remarkably low mutational burden [20]–[22]. Deficiency in SMARCB1, a component of the SWI/SNF chromatin remodeler complex, leads to altered genome-wide DNA methylation, dysregulated epigenetic control, and aberrant gene expression signatures [23]–[25]. Based on DNA methylation and gene expression signatures, distinction of three major ATRT subtypes, termed ATRT-MYC,-TYR and-SHH has been established [26]. The subtypes show distinct anatomical sites at time of diagnosis and likely reflect different cellular tumor origins [3], [26]. ATRT-TYR tumors have been associated with better overall survival [27], [28]. Further, ATRT-SHH have been proposed to split up into two subtypes based on DNA-methylation, though this distinction carries no prognostic difference [29]. Despite many efforts for targeted therapeutic interventions, treatment of ATRTs is currently limited to high precision surgery, radiotherapy and chemotherapy with a dismal 3-year overall survival rate of 25% [30], [31].

The successful application of immunotherapy has been a therapeutic revelation for many tumor entities, with a focus on tumors with high neoantigen load, such as DNA mismatch repair deficient tumors or tumors with high mutational burden [32] (as well as clinical trials: NCT02912559, NCT02563002, NCT02983578, NCT02992964, NCT03040791 and NCT02359565). As SWI/SNF-deficient tumors, ATRTs are characterized by a simple genome with low tumor mutational burden, and thus have been considered poor sources for tumor-mutated neoantigens. However, a distinct immune infiltration has been described in ATRT subtypes and studies showed that the microenvironment of ATRT is more immunologically active as compared to other pediatric CNS tumors [33]–[36]. Hence, ATRTs might be targetable by immunotherapy (e.g. utilizing immune checkpoint blockade) [37], [38]. Leruste et al. showed that ATRTs are infiltrated by immune cells, including clonal T cells, indicating a tumor-specific response [37]. Instead of mutation-driven immunogenicity, the study provides evidence that transcriptional activity of classically non-protein-coding, repetitive genomic sequences of the transposable element (TE) type drives ATRT immunogenicity. Specifically, the authors show that sense and anti-sense transcription of endogenous retroviruses correlates with cytolytic score and reason that consequential double-stranded RNA-mediated sensing and type I/III interferon response are responsible for inducing ATRT immunogenicity [37]. This study first highlighted a potential role for TEs in ATRT tumor biology.

TEs make up around 50% of the human genome content and are grouped into two main classes based on their original mode of propagation [39], [40]. Class I elements, termed retrotransposons, propagate by a *copy&paste* mechanism via an RNA intermediate. Conversely, class II elements, termed transposons, replicate by a *cut&paste* mechanism. Within class I elements, Long Terminal Repeat (LTRs) and Long Interspersed Nuclear Elements (LINEs) outnumber other elements by far, constituting as much as 8% and 20% of the human genome, respectively [40]–[42]. Pioneering computational work on these repetitive sequences allowed further classification into families and subfamilies, providing information on evolutionary ancestry and their exact genomic positioning within the human reference genome (UCSC RepeatMasker track, hg38) [4], [43]. In the past decade, there has been a paradigm shift in our understanding of the role of LTR and LINE1 elements in human health and disease. Classically, TEs have been considered as degenerate, non-functional genomic elements that underlie tightly repressed through epigenetic silencing in adult cells and tissues [44]. This conception has shifted with increasing evidence of evolutionary co-option of TEs, at the level of transcripts, proteins, or cis-regulatory genomic elements [44]–[46]. In the context of cancer, a number of studies demonstrate that TE transcriptional control is at least partially lost, resulting in increased LTR and LINE1 transcriptional activity as part of global gene expression changes [47]–[52]. In general, oncogenic transformation appears to coincide with transcriptional activation of LINE1 and LTR elements. This activation has a dichotomous effect on the cell. On one side, it promotes tumor growth – for example, through aberrantly active TE-derived regulatory elements that drive expression of oncogenes [53]. On the other hand, it has been shown, that the altered transcriptome profile of LTR elements can also induce an anti-tumor response through innate immune activation, as suggested for ATRTs [54], [55].

Here, we investigate whether transcriptional dysregulation of LTR and LINE1 elements contributes to the immunogenic landscape and molecular identity of ATRTs. Using TE-focused transcriptomic profiling of primary tumor samples from all three ATRT subtypes, we investigate the expression of 1.9M LTR and LINE1 elements at family and locus-level resolution. We integrate our data with expression profiles of protein-coding genes and undertake comparative profiling in ATRT cell line models, often used to study tumor biology and therapeutic interventions. Our data demonstrate that ATRTs possess a subtype-specific signature of TE activity, wherein ATRT-MYC tumors stand out with particularly low LINE1 activity. Given the standing need for optimized therapeutic regimens to tackle this aggressive pediatric tumor entity, our data provide an important resource for future studies that focus on understanding ATRT molecular identity and promoting potential TE-driven ATRT immunogenicity.

## Results

### 1. TE transcriptome profiles define the identity of ATRT subtypes

We set out to determine the TE transcription profiles in ATRT-MYC,-TYR and-SHH tumors at subfamily and locus-level resolution. Our analysis is based on a set of primary human ATRT samples [2], [3], that had been stratified for tumor subtype by their DNA methylation pattern using the DKFZ brain tumor methylation-based classification [56]. In Fig. 1a-c we present the clinical information about the CNS anatomic location of the primary tumor diagnosis based on magnetic resonance imaging (MRI), sex and age along with additional meta-data of the classifier score (Classifier version 12.8) and ATRT subgroup. We have obtained samples with matching transcriptome and DNA methylation data (n=10) for further analysis [2], [3]. RNAseq samples were processed for locus-specific analysis of TE transcripts using the SalmonTE computational pipeline. This sample subset comprised all three ATRT subtypes (2x TYR, 4x MYC, 4x SHH). We used the RepeatMasker (hg38) annotation of TEs, and focused our analysis on LINE1 and LTR entities as the main source of TE-derived transcripts. LTR elements are further classified into ERV1, ERVL, ERVK and ERVL-MaLR families. In addition, LTR and LINE1s, comprise several hundred subfamilies, which are summarized in Supplementary Table 1. Overall, our annotation includes 792,000 LTR and 1.19M LINE1 genomic elements.

**Fig. 1:**
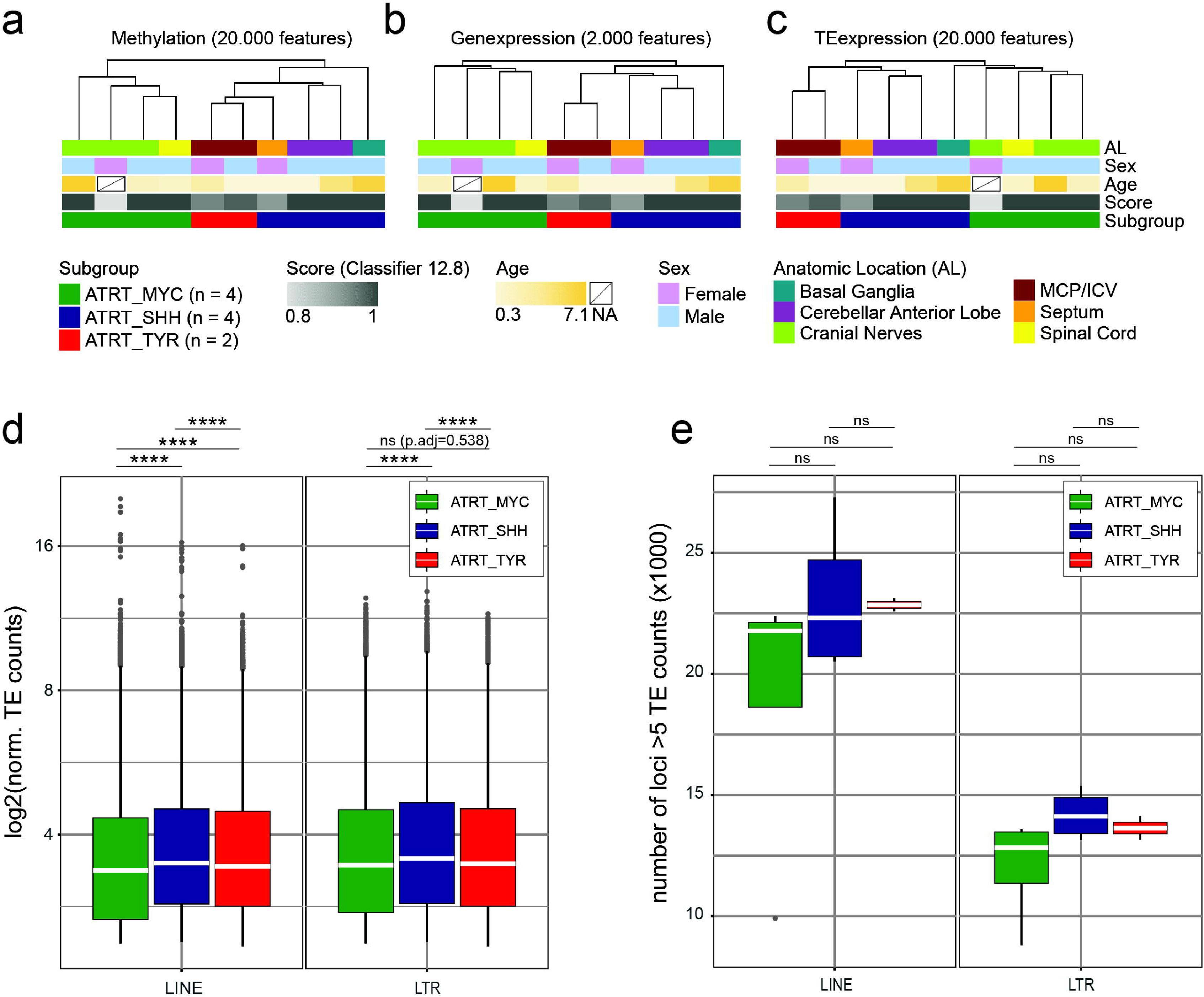
Primary samples show ATRT subtype clustering based on TE transcription profiles. Unsupervised hierarchical clustering of a) DNA methylome beta values (20,000 most variable CpGs), b) gene transcription (2,000 most variable genes) and c) TE transcription (20,000 most variable transcripts). ATRT subgroup classification based on DNA methylome analysis by Capper et al., 2018 [56]. d) DESeq2 normalized read counts from loci with at least 5 mapped reads. Median read count is indicated as white bar. e) Number of expressed TE loci with at least 5 mapped reads. Statistical analysis one-way ANOVA with post-hoc Tukey’s Honest Significant Differences (HSD) test. **** p < 0.00001.

As a first step, we validated the datasets based on DNA methylation array and gene transcriptome analysis. Regarding the DNA methylome, unsupervised hierarchical clustering based on top 20,000 most variable beta values confirmed grouping of the samples into the established ATRT subtypes (Fig 1a). Regarding their overall gene transcriptome profile, the samples showed unsupervised hierarchical clustering into ATRT subtypes analogous to their methylome profile (Fig 1b).

To investigate gene transcriptomic signature in further detail, we undertook differential gene expression (DEG) analysis between the samples of established subtypes. DEG analysis yielded ATRT subtype-specific gene expression (Suppl. Fig. 1a), with 223 differentially expressed genes across all three ATRT subtypes (MYC_DEG_ = 69, SHH_DEG_ = 65, TYR_DEG_ = 89). We particularly observed a set of subtype-specific genes (MYC/TYR/SHH-cluster-genes) that were transcriptionally active specifically in one ATRT subtype. We cross-referenced these results to 42 previously-described ATRT subtype differences in gene transcriptional activity and observed that the transcriptomic datasets reliably reflected these differences (Suppl. Fig. 1b) [23], [24], [26], [57]–[61].

In summary, these results show validity of described transcriptomic datasets for our analysis. We therefore proceeded to assess genome-wide transcriptional activity of TEs. Regarding overall TE transcriptome profiles, we observed ATRT subtype specific clustering that mirrored DNA methylation and gene transcription-based analysis (Fig 1c). ATRT subtypes can thus be defined by the genome-wide signature of TE transcriptional activity analogous to profiling by DNA methylation or gene transcription data (Fig 1a-c).

TEs are distributed across the entire genome, with 43% of elements positioned within gene bodies and around 25% positioned at over 50kb distance of the next gene transcriptional start site (TSS) (Table 1). We asked if positioning of TEs relative to genes could affect the observed TE-transcriptome based ATRT subtype specific signature. We therefore grouped all elements into TEs within genes (43.6%), TEs up to 10 kb upstream of an annotated transcriptional start site (TSS) (7.1%), TEs between 10 to 50 kb upstream of a TSS (24.2%) and TEs >50 kb upstream of a TSS (25.2%) (Suppl. Fig 2a, Table 1) and performed principal component analysis (PCA) for each TE subset. PCA based on any of our defined TE subsets also showed a consistent separation of the ATRT subtypes along the first two principal component vectors, reflecting PCA based on all TE or gene transcripts (Suppl. Fig 2a). The ATRT TE transcriptome thus reflects the tumor subtype-specific functional genome faithfully independent of the genomic position of the TE entities.

**Table 1.** Genomic location in relation of gene transcriptional start sites.

| **genomic region** | **# of elements** | **%** |
| --- | --- | --- |
| within genes | 1.06M | 43.6 |
| 10kb upstream TSS | 172K | 7.1 |
| 10-50kb upstream TSS | 589K | 24.2 |
| >50kb upstream TSS | 612K | 25.2 |

We next went on to analyze the ATRT TE transcriptome in further detail, focusing on differences in total TE-derived transcript counts and number of transcribed TE loci between ATRT subtypes. Notably, the SalmonTE pipeline aligns reads to TE coding strand and thus is not skewed by anti-sense read mapping. To minimize the impact of loci with low number of mapped reads in our analysis we considered TE loci with a normalized read count of at least 5 (Fig. 1d, e) [62]. Results were stratified by TE class, namely LINE1s and LTRs. ATRT-MYC subtype showed the lowest transcriptional activity on average both for LINE1s (log2(median±SD): MYC 3.37±1.3, SHH 3.48±1.35, TYR 3.43±1.34) and LTR elements (log2(median±SD): MYC 3.45±1.38, SHH 3.56±1.38, TYR 3.47±1.35)

(Fig. 1d). Similarly, ATRT-MYC tumors showed a trend towards the least amount of actively expressed loci again both for LINEs (median±SD: MYC 21,778±6,047; SHH 22,316±3,170; TYR 22,855±384) and LTR elements (median±SD: MYC 12,821±2,229; SHH 14,113±1,048; TYR 13,636±689) (Fig. 1e). Taken together, ATRT-MYC subtypes are characterized by significantly less LINE1 and LTR transcripts that tend to be derived from fewer TE loci.

We next went on to define expression of LTR and LINE1 elements at subfamily level in the three ATRT subtypes (Fig. 2). Our analysis focused on TE subfamilies with at least 5 aligned reads on average in at least one ATRT subtype and we plotted data according to expression level (relative mean of all expressed elements per subfamily) and percentage of expressed TE loci per subfamily (occurrence score). We separated the LTR class into the four LTR families ERVK, ERV1, ERVL and ERVL-MaLR (Fig. 2 ERVK family, Suppl. Fig 3 ERV1, ERVL and ERVL-MaLR). Our analysis reveals that ATRT subtypes differ considerably in LTR subfamily expression. For example, while LTR5B (ERVK) elements show strongest expression in ATRT-TYR, LTR5A (ERVK) express highest in ATRT-MYC (Fig. 2A). In contrast, LINE1 and LTR ERVL-MaLR subfamilies showed consistently lower expression in ATRT-MYC tumors compared to ATRT-SHH and-TYR subtypes for most LINE1 subfamilies (Fig. 2B, Suppl. Fig 3). Only few subfamilies, e.g. MARE6 or LINE1MCa, showed an opposing trend, with highest expression in ATRT-MYC tumors. Occurrence scores ranged from 0.3 to 6 % across all subfamilies, indicating that individual elements are transcriptionally active rather than entire subfamilies.

**Fig. 2:**
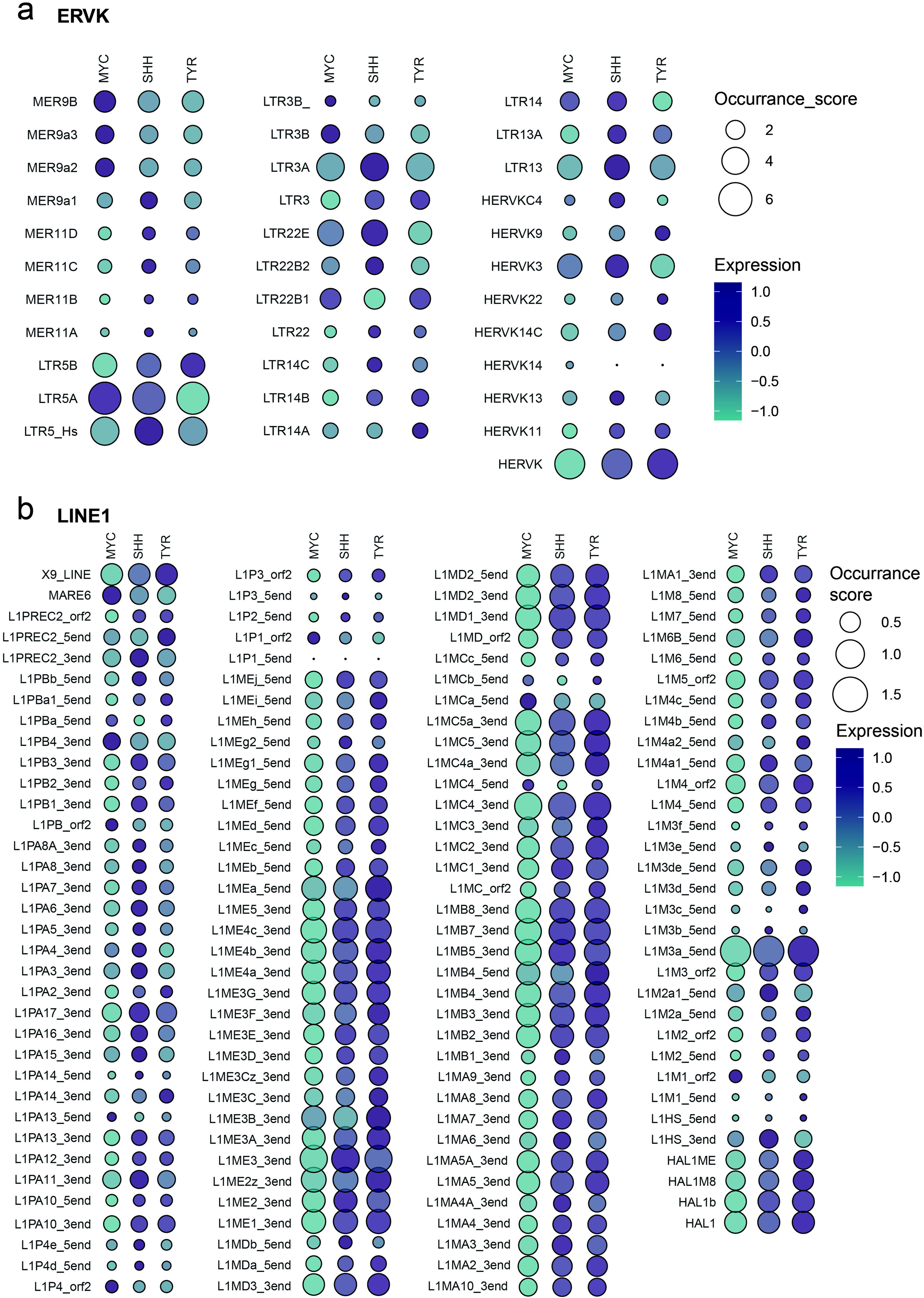
Differences in TE transcription at subfamily level Relative normalized TE counts were summarized at TE subfamily level, including elements with at least 5 mapped reads on average in at least one ATRT subtype (expression score). Occurrence score is defined as the number of expressed loci per subfamily [in %]. a) Subfamilies of the ERVK family (LTR elements) (other LTR families presented in SupplFig 3). b) Subfamilies of the LINE1 family.

In summary, our analyses demonstrate that ATRTs show a subtype characteristic TE transcription signature that likely reflects tumor subtype-specific transcriptional regulation of TEs. ATRT-MYC tumors show overall the lowest counts of TE-derived transcripts and the lowest number of transcribed TE loci, with in particular reduced transcriptional activity of LINE1 subfamilies in comparison to ATRT-SHH and ATRT-TYR subtypes. ATRT identity is thus defined not only by a characteristic gene transcriptional program, but also by characteristic activity of LTR and LINE1 genomic elements.

### 2. ATRT subtypes are marked by specifically transcribed TE loci

We next focused our analysis to locus-level resolution and identified differentially expressed (DE) LTR and LINE1 elements by pair-wise comparison of the ATRT subtypes. The identified DE TEs are presented in the heatmaps shown in Fig. 3a and b. We detected a total of 383 and 466 differentially expressed LTR and LINE1 loci, respectively (cutoffs log2FC>|0.5|, p_adj_<0.05). Each ATRT subtype showed transcription of a characteristic cluster of LTR elements (MYC_LTR_ 34; SHH_LTR_ 175, TYR_LTR_ 174) and LINE1 elements (MYC_LINE1_; SHH _LINE1_ 190, TYR _LINE1_ 212), which we termed MYC-, SHH-and TYR-‘cluster.TEs’ (Fig. 3a,b). These subtype-specific cluster elements showed different composition with regard to LTR families (Fig. 3c) and LTR/LINE1 subfamilies (Fig 3d, e). Statistical dependence analysis between ATRT-cluster.TEs and LTR families revealed significant results (χ^2^ = 17.533, Fisher p-value=0.0075) and highlighted in particular ERVL-MaLR elements being overrepresented in MYC and SHH clusters, as well as ERV1 elements in the TYR cluster.

**Fig. 3:**
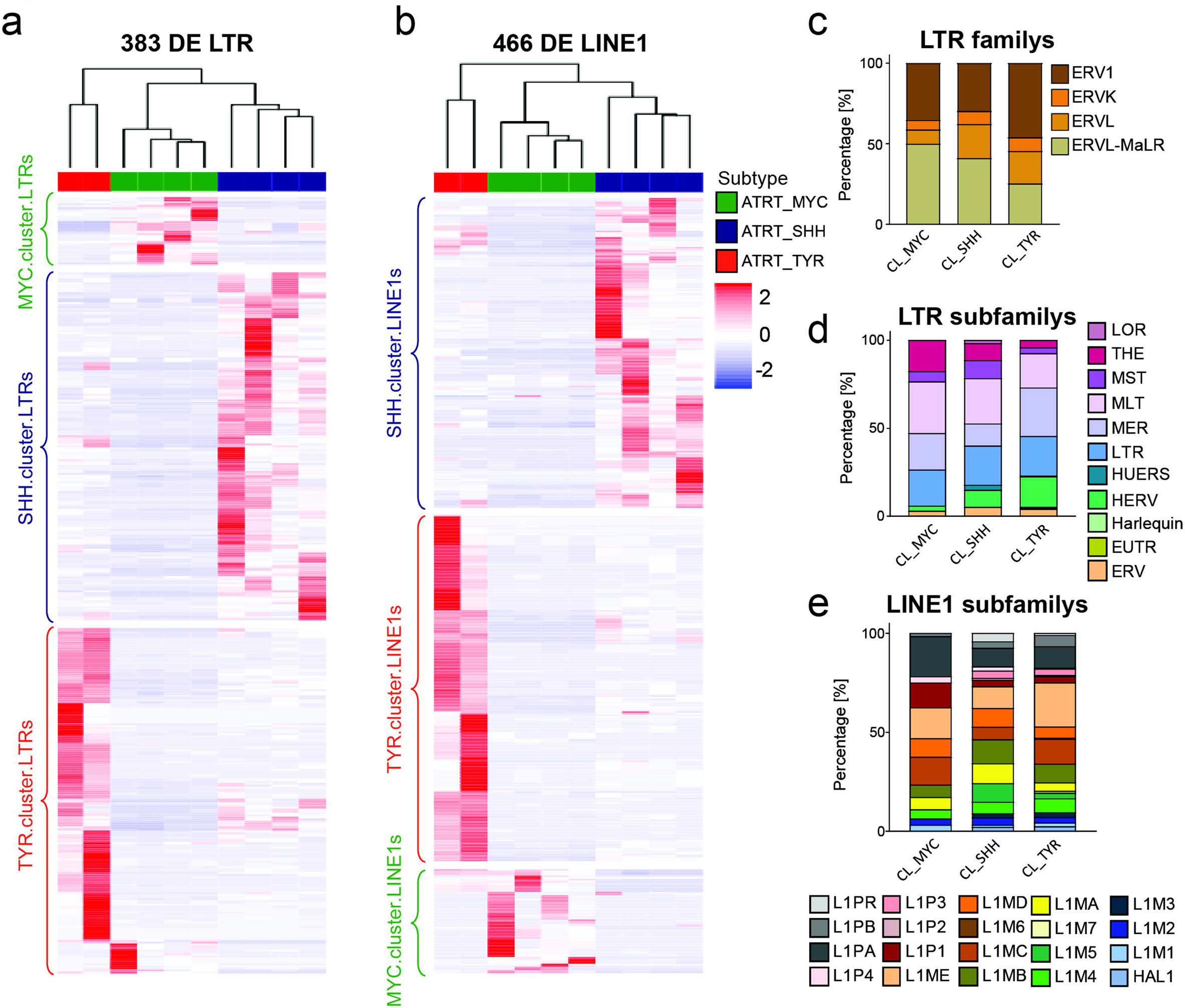
Differential TE transcription analysis Differential LTR (a) and LINE1s (b) as determined by DESeq2 with cutoff log2FC>|1| and p-adj <0.05. Unsupervised hierarchical clustering of samples and elements reveals ATRT subtype-specific differentially transcribed LTRs and LINE1s, respectively (termed ‘cluster.TEs’). Relative fractions of clusterTEs at LTR family c) and subfamily level d). For better visualization LTR subfamilies are grouped based on their annotation name. e) Relative fractions of clusterTEs for LINE1 subfamilies.

In order to assess whether cluster.TEs represent selected TEs regarding their genomic position in relation to human genes, we grouped TEs in bins with increasing genomic distance to the closest gene transcriptional start sites (bins identical as presented in Suppl. Fig 2a) and compared the frequency of cluster.TEs to the number of all transcribed elements (including non-differentially expressed loci). We observed for all subtype clusters that LTR elements residing within genes were over-represented in ‘cluster.TEs’ compared to their frequency in all transcribed TEs (Fig. 4a). On the other hand, differential transcribed LTRs elements residing >50 kb upstream of a TSS were consistently under-represented in all three ATRT clusters (Fig. 4a). Differentially expressed LINE1s showed a different profile in the context of genomic location (Fig. 4b). Gene-and proximal-to-gene residing LINE1 elements of the MYC-cluster were consistently under-represented. In contrast, distal LINE1 elements (>50 kb distance to TSS) were over-represented in MYC-cluster.LINE1s, reflecting a marked distinction of TE transcription profiles of ATRT-MYCs compared to ATRT-SHH and –TYR. Taken together, each ATRT subtype is defined by an individual transcriptional signature of TE loci (‘cluster.TEs’).

**Fig. 4:**
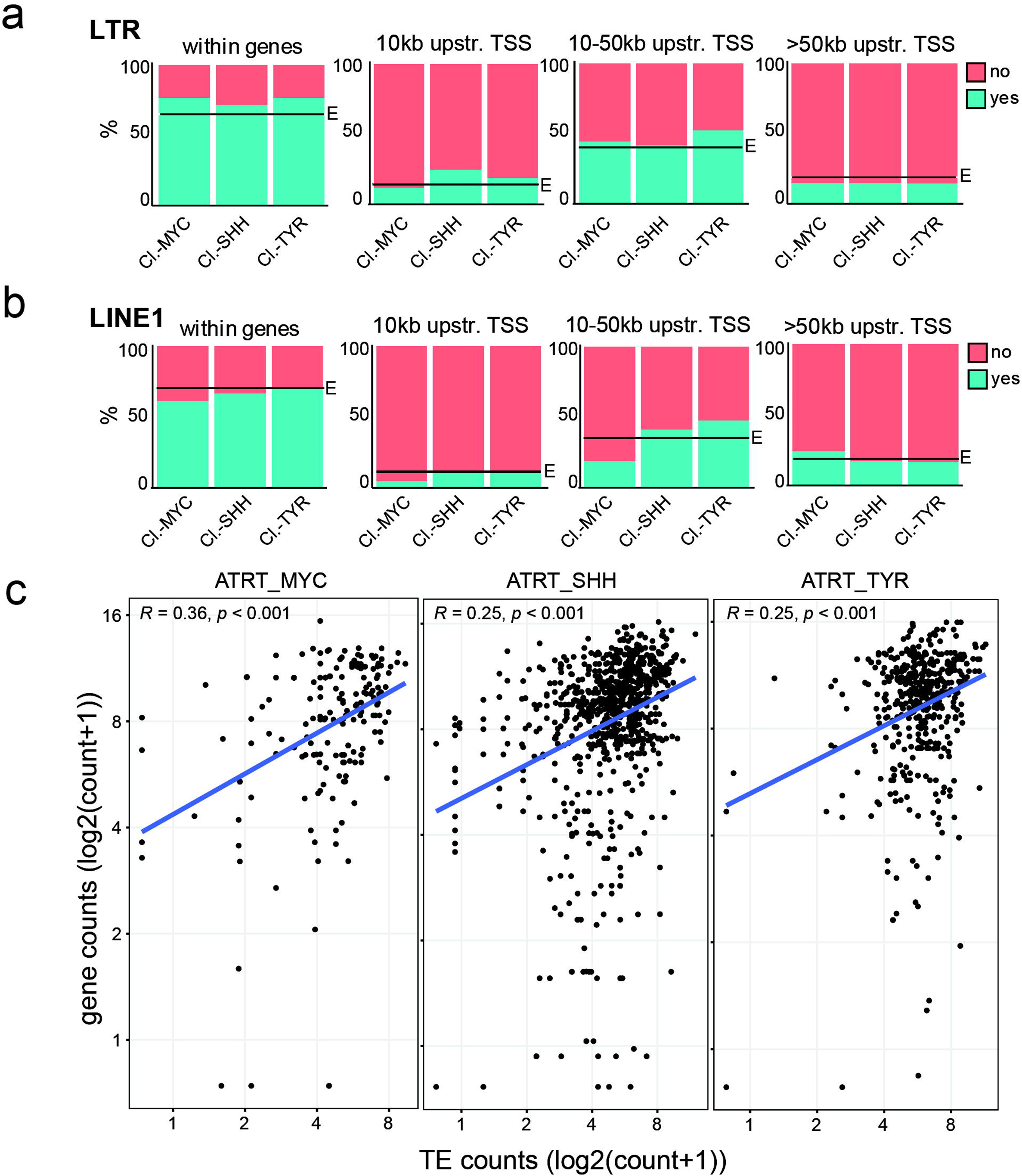
Active TEs in context of genomic distance to gene transcriptional start sites Active clusterTEs in relation to their genomic distance to gene transcriptional start sites for LTRs a) and LINE1s b), respectively. Actively transcribed elements = “yes” (turquoise), not transcribed = “no” (red). “E” indicates the observed frequency taking all transcribed LTR/LINE1s into account. c) Correlation analysis of gene and TE transcription for ATRT.clusterTEs residing within gene bodies. DESeq2 normalized counts, excluding TEs and genes with zero counts. Pearson correlation and p-value are indicated.

About half of the transposable elements reside within genes (in introns) and 85-90% of all human genes contain TEs [63]. Thus, elevated TE transcription could be a consequence of increased gene expression. To investigate this, we correlated the transcription level of ‘cluster.TEs’ with the transcription level of the genes they reside in. We find statistically significant correlation for all three ATRT subtypes and their respective cluster TEs (Fig 4c), indicating that gene transcription can influence TE transcription and vice versa.

### 4. ATRT cell line models partially reflect the TE transcription signature of primary ATRTs

Mechanistic studies on ATRTs often rely on ATRT-derived cell line models. These cell lines have been associated to ATRT subtypes based on DNA methylome and transcriptome data, which show good correlation when compared to each other [26]. We therefore asked whether observed TE transcript signatures in primary ATRT samples are reflected in ATRT cell lines. Since ATRT-TYR generally lacks established cell line models, we focused on BT12, BT16 and CHLA06 cell lines for ATRT-MYC-type and CHLA02 cells as an ATRT-SHH-type model (cell line association with tumor subtype based on literature) [26], [64]. Deep transcriptomic samples were generated in triplicates from each cell line and submitted to Salmon and SalmonTE for read alignment. PCA based on all gene transcripts showed high diversity between cell lines and no clear grouping according to tumor subtype (Fig. 5a). PCA based on TE transcripts mirrored this result (Fig. 5a). Notably, as seen for primary ATRT samples, TE transcriptome profiles in PCA resemble gene transcriptome profiles, supporting the notion that TE transcriptomes are tumor-type specific (see Fig. 1a-c Suppl. Fig. 2b). As for the quantity of TE transcripts, cell models show comparable levels to primary ATRT samples (average log2(median) transcripts 3.46 and 3.33 in primary and cell line data, respectively). However, TE transcripts in cell line samples are derived from fewer loci compared to primary data (average loci number 17,920 and 11,892 in primary and cell line datasets, respectively). Among cell line datasets, ATRT-MYC shows significantly lower LINE1 transcript levels compared to ATRT-SHH (log2(median±SD): MYC 3.25±1.23; SHH 3.27±1.13), consistent with our analysis in primary samples (Fig. 5b). Interestingly the opposite trend applies for LTR transcripts. Here, ATRT-MYC cell line samples show more transcripts compared to ATRT-SHH (log2(median±SD): MYC 3.40±1.39; SHH 3.39±1.22), which is reflected across the three LTR families ERV1, ERVL and ERVL-MaLR (Fig. 5b). Regarding the number of expressed TE loci, cell lines are consistent with primary data in that ATRT-MYC shows a trend in expressing fewer LINE1 and LTR loci compared to ATRT-SHH (median±SD; LINE1: MYC 13,599±2,102; SHH 15,506±2,507; LTR: MYC 9,026±1,141; SHH 9,437±944) (Fig 5c).

**Fig. 5:**
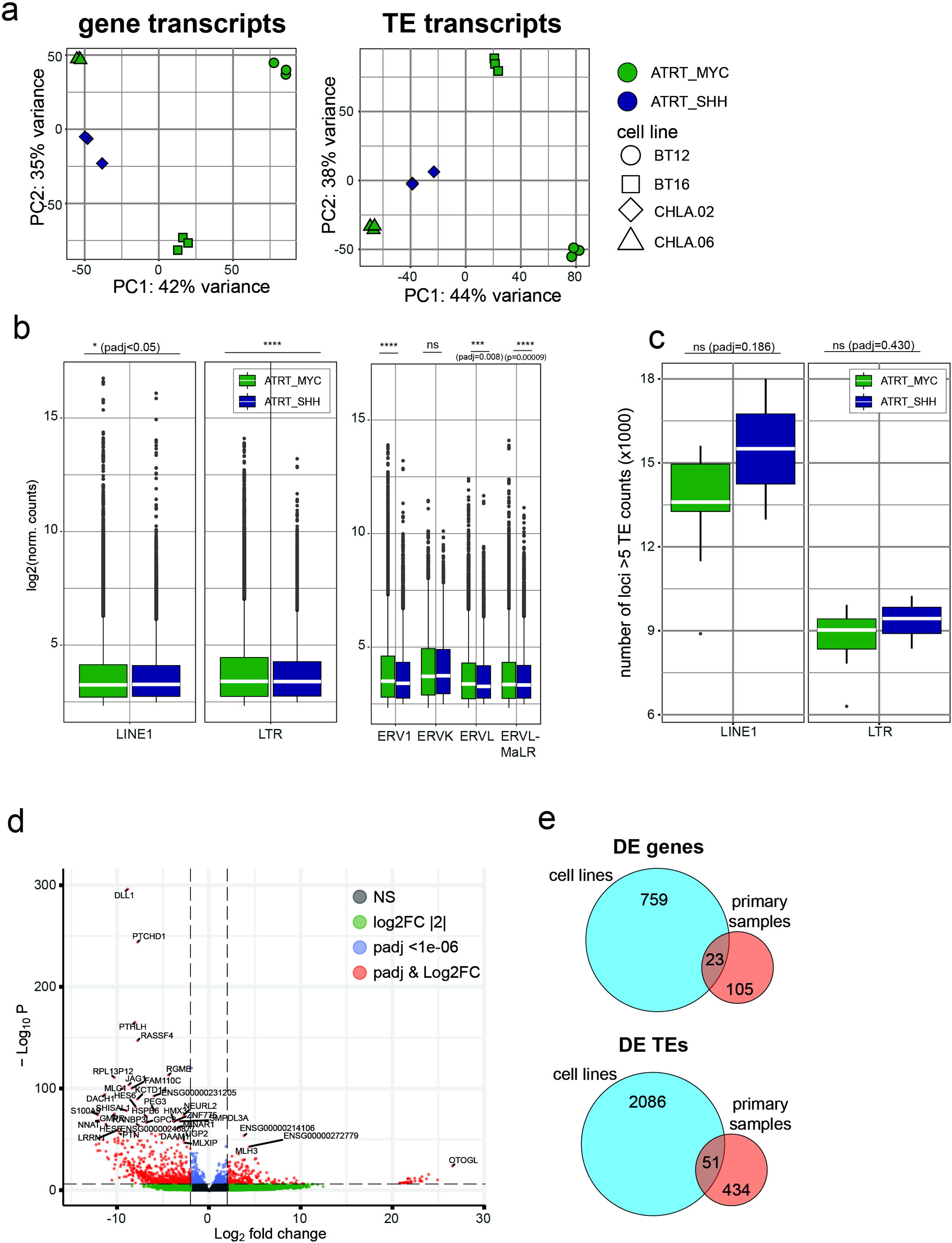
TE transcriptional profiles in established cell models of ATRT-MYC and-SHH Comprehensive LTR and LINE1 transcriptional profiling for three ATRT-MYC cell lines (BT12, BT16, CHLA.06; n=3 each) and ATRT-SHH cell line CHLA.02 (n=3). Cell line assignment to ATRT tumor subtype based on literature consensus. a) Principal component plot for 500 most variable gene and TE transcripts, respectively. b) Overall transcript abundance for TE classes LINE1 and LTRs, as well as LTR families ERV1, ERVK, ERVL and ERVL-MaLR. Elements with at least 5 transcript counts included. Median transcript level indicated by white bar. c) Number of active LINE1 and LTR loci, respectively. Loci with at least 5 transcript counts included. Median indicated by white bar. Statistical analysis in b) & c) one-way ANOVA with post-hoc Tukey’s Honest Significant Differences (HSD) test. **** p < 0.001, *** p < 0.01, * p < 0.05. d) Differentially expressed genes contrasting ATRT-MYC vs. ATRT-SHH. Cutoff thresholds log2FC>|2|, padj < 1×10E-06. e) Number of differentially expressed genes and differentially transcribed TEs in cell lines vs primary samples and their respective overlap. Thresholds were set at log2FC>|2|, padj < 1×10E-06 for DEGs and log2FC>|1|, padj < 0.05 for DE TEs.

Differential gene expression analysis between MYC and SHH cell line models revealed 219 up-and 564 downregulated genes (log2FC>|2|, p<10E-06) (Fig 5d) and 103/107 up-and downregulated TEs, respectively. In general, cell line datasets yielded more differentially expressed genes (782 vs 128) and TEs (2137 vs 485) compared to primary data, with a fractional overlap between both sample sets (23 shared DEGs and 51 shared TEs between cell lines and primary samples) (Fig. 5e) (Information on shared DEGs and TEs provided in Supplementary Table 2 and 3, respectively). Taken together, these results show that ATRT cell line models, in particular for ATRT-MYC subtypes are heterogenous regarding their overall gene and TE transcriptome. Furthermore, they only partly reflect transcriptomic subtype differences of primary ATRT samples.

We next asked, to what extent ATRT-MYC and-SHH cell lines reflect the subtype-specific ‘cluster.TE’ signature, as defined by our analysis of primary ATRT sample datasets (refer to Fig. 3a, b). PCA combining both primary and cell line datasets revealed co-clustering of both dataset entities according to their ATRT subtype considering the “sample type” parameter as a co-variate in batch correction of the data (Fig. 6a, b). This indicates that distinct differences in gene and TE transcription profiles exist, allowing differentiation into ATRT subtypes. Next, we focused on all MYC and SHH defining ‘cluster.TEs’ from primary ATRT samples (see Fig. 3). We found 20 LTR and 43 LINE1 elements from the MYC.TE cluster and 157 LTR and 140 LINE1 elements from the SHH.TE cluster to be expressed in the cell lines. We then plotted their cell line expression values using unsupervised hierarchical clustering (Fig. 6c, d). This showed a clear clustering of cell line samples according to tumor subtype. In detail, we found that 70% of SHH-defining cluster.LTRs (53% of cluster.LINE1) show consistent transcription in the SHH cell line, and 65% of MYC-defining cluster.LTRs (70% of cluster.LINE1s) are consistently transcribed in the MYC cell lines (χ^2^ test: LTRs χ^2^=8.632, *P*=0.003; LINEs χ^2^=5.875, *P*=0.015). Taken together, these data demonstrate that the subtype-defining TE expression signatures of primary ATRT samples are partially reflected in ATRT cell line models, allowing for use of these models in future studies that focus on TE activity in ATRT biology.

**Fig. 6:**
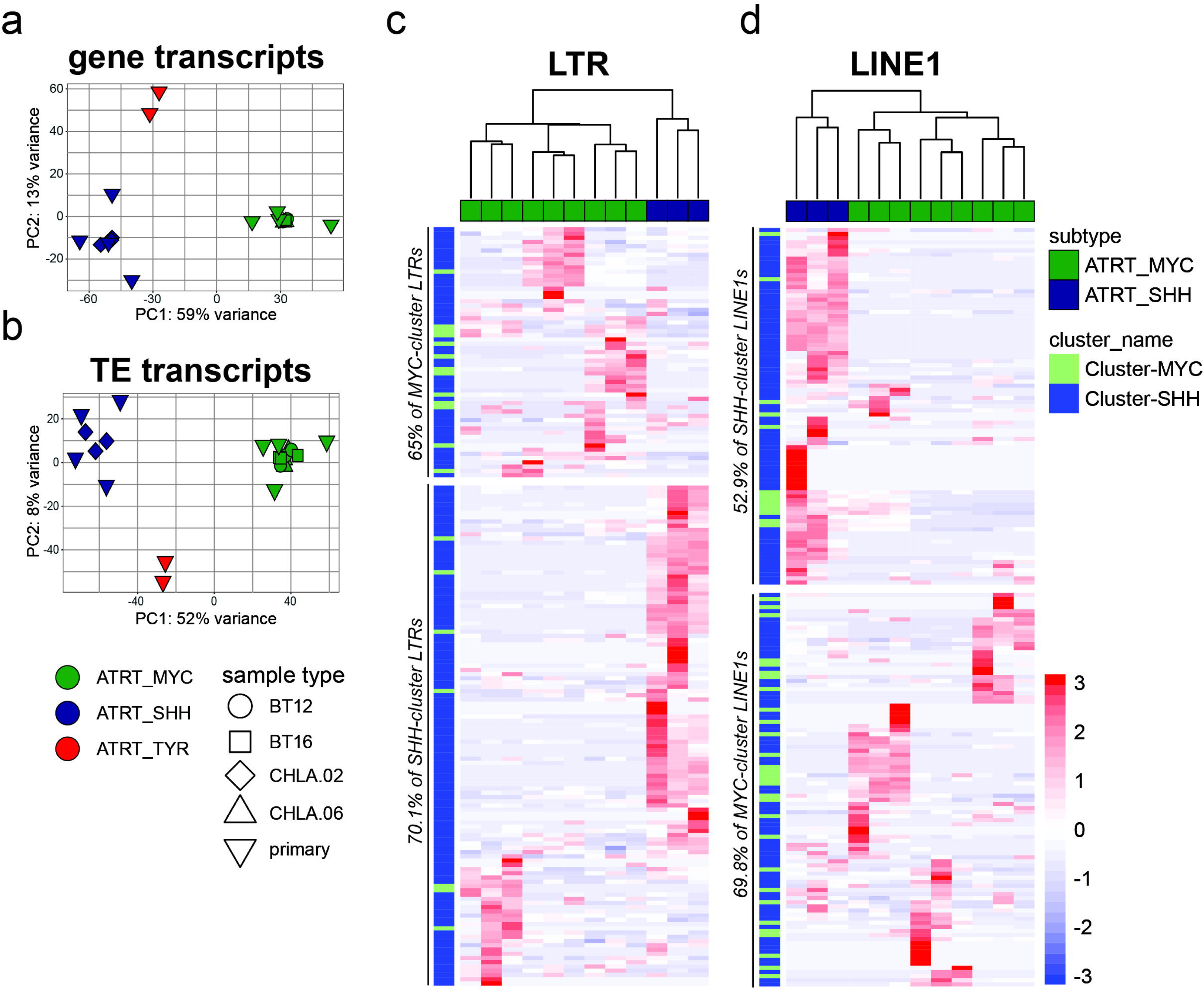
Combined analysis of primary and cell model transcriptome data a) and b) Principal component plot for 500 most variable gene and TE transcripts integrating data with co-variate correction for “sample type”. Unsupervised hierarchical clustering of MYC-and SHH-clusterTEs found transcribed in the cell line data for LTRs c) and LINE1s d).

### 5. TE transcription in ATRT subtypes as potential driver for tumor immunogenicity

Expression of TEs has been associated with tumor immunogenicity through TE-derived promoter activity, TE peptide expression or activation of innate immune responses through TE transcripts [65]–[67]. In ATRTs, bidirectional transcription of LTR elements has been suggested to drive interferon responses through double-stranded RNA-mediated sensing in MYC subtypes [37]. We therefore set out to investigate whether, TE transcriptome-associated immunogenicity features differ across ATRT subtypes. We first studied bidirectional expression levels of LTRs and LINE1s in primary ATRT samples. To do so, we processed transcriptomics datasets for TE transcript counts in the sense and antisense direction at locus-level resolution. All TE loci with at least 10 read counts in both sense and antisense direction in at least 50% of the samples from one ATRT subtype were defined as bidirectionally transcribed for that subtype. We observed significant differences in bidirectionally transcribed TE loci between ATRT subtypes: As already observed for sense transcription (see Fig. 1e), ATRT-MYC tumors show the lowest number of active LINE1s and LTR loci with bidirectional activity, while ATRT-TYR samples show the highest number (Fig. 7a, b). Interestingly, the overall levels of bidirectional TE-derived transcripts are similar between all three ATRT subtypes. However, ATRT-MYC tumors show a tendency for increased levels of LTR (Fig. 7c) and decreased levels of LINE1s bidirectional transcripts (Fig. 7d), compared to ATRT-SHH and-TYR cases. Thus, while tumors of the ATRT-MYC subtype have fewer active LTR loci that show bidirectional transcription, these loci appear to have higher transcriptional activity compared to other subtypes.

**Fig. 7:**
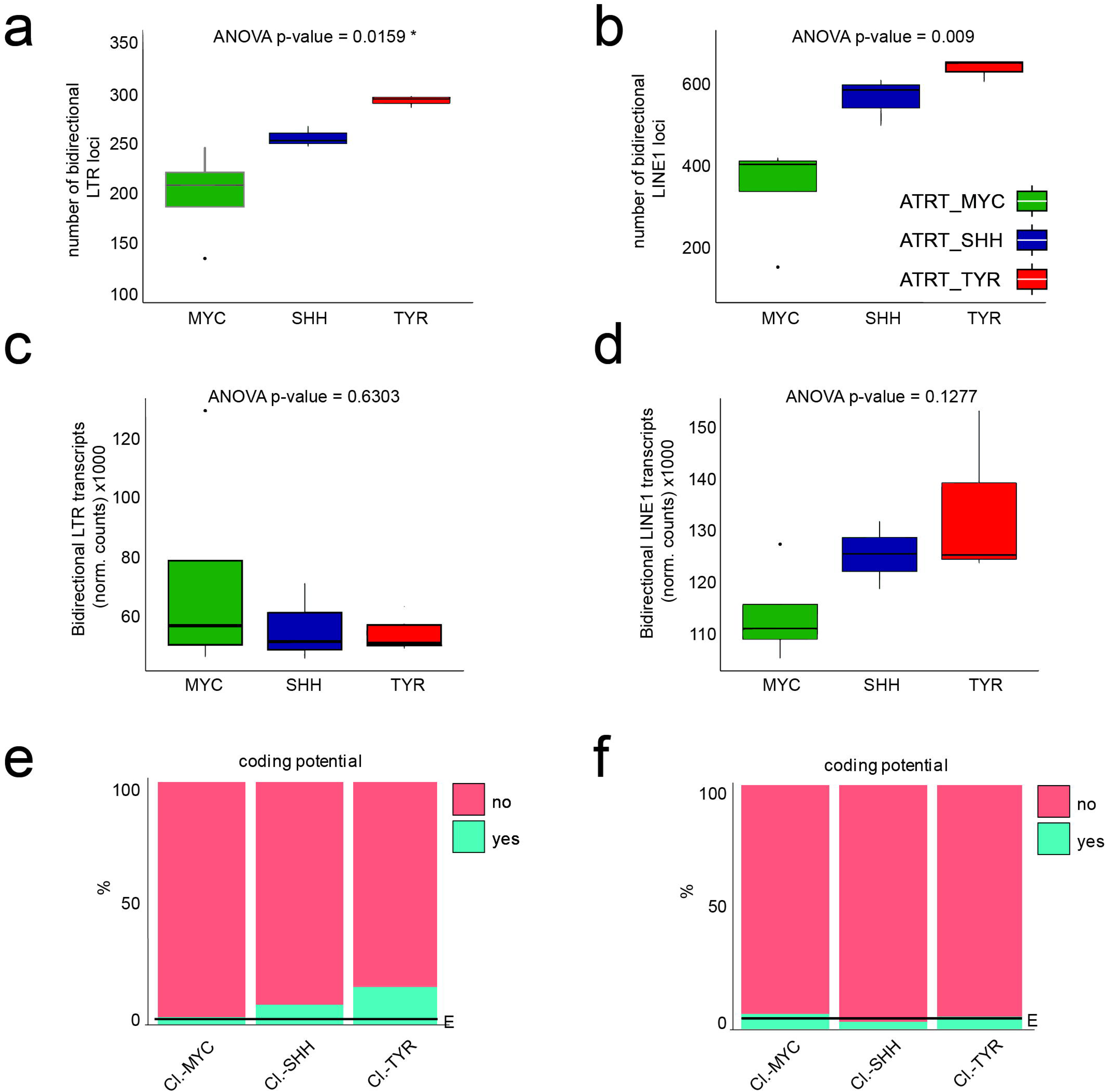
Bidirectional transcription from LTRs and LINE1s in ATRT subtypes and protein-coding potential a) and b) number of LTR and LINE1 loci in ATRT subtypes with bidirectional transcriptional activity. c) and d) transcript levels of LTR and LINE1 loci with bidirectional transcription. Statistical test one-way ANOVA. Active clusterTEs with protein-coding potential according to literature for LTRs e) and LINE1s F), respectively [68]. Actively transcribed elements = “yes” (turquoise), not transcribed = “no” (red). “E” indicates the observed frequency taking all transcribed LTR/LINE1s into account

Since TE-derived peptides, as tumor-specific antigens, have also been associated with tumor immunogenicity, we secondly investigated the expression levels of potentially protein-coding TEs in the primary ATRT samples. LTR and LINE1 loci with putative open reading frames for peptide or protein expression were selected based on a published dataset and cross-referenced to the TE annotation that was used for transcriptome analysis [68]. We observe a prominent over-representation of potentially protein-coding LTR elements in ATRT-SHH and-TYR subtypes (Fig. 7e) and a minimal over-representation of potentially protein-coding LINE1 elements in ATRT-MYC (Fig. 7f) in relation to all expressed TEs, indicating another differential characteristic between ATRT subtypes with potential implications for effective immunotherapy.

## Discussion

Studies towards an improved understanding of the gene regulatory network of ATRTs have been driven by the need to develop advanced therapeutic strategies for this hard-to-treat tumor entity with devastating prognosis. In this context, recent landmark studies have focused on characterizing the molecular diversity of ATRT subtypes. These studies have highlighted, that diversity is likely rooted in different lineages of origin for MYC, TYR and SHH subtypes [3]. Furthermore, they have suggested that differences in genome-wide transcriptional landscapes could render certain ATRT tumors immunogenic and thus responsive to immunotherapy [26], [31], [37]. This phenotype was suggested to be linked to the transcriptional activity of TEs, which appear to be dysregulated in ATRTs as a result of *SMARCB1* deficiency and associated genome-wide epigenetic remodeling [37], [69]. Lastly, expression of HML2 LTR elements has been suggested as putative driver for tumorigenesis in ATRTs [69]. These studies demonstrate the importance to first, study and understand ATRT subtype diversity in further detail, and to second, investigate transposable element activity in ATRTs.

We here set out to address these points. Using a computational pipeline for transcriptome analysis of repetitive elements at locus-level resolution, we present the first comprehensive TE expression profiling of primary ATRTs across MYC, TYR and SHH subtypes. Our transcriptome analysis comprises 1,9M TEs, with a particular focus on LTR and LINE1 elements. These two classes of TEs account for over 25% of human genomic sequences and their activity has been implicated in mediating interferon response and neoantigen generation in ATRTs and cancer cells in general [38], [65], [70], [71]. Our data show that ATRT subtypes are defined by a genome-wide signature of LTR and LINE1 element transcriptional activity. This signature allows to identify subtypes similar to classification based on DNA methylation or gene transcriptome. We show expression differences at TE family level and also define a set of 383 LTR and 466 LINE1 elements, here termed ‘cluster.TEs’, that show specific expression in one ATRT subtype, respectively. Transcriptional activity of these cluster TEs also allows subtype differentiation of immortalized ATRT cell line models that generally are transcriptionally heterogeneous.

Since TEs, as human genes, underlie definite epigenetic control, our study suggests subtype-specific epigenome alterations that affect gene and TE activity. This raises the question of a potential crosstalk between TE and gene transcriptional activity. Indeed, TEs and in particular LTR elements, have been shown to act as gene promoters and enhancers in different cellular contexts and approximately 25% of human cis-regulatory elements are located within LTRs [52], [71]–[75]. We observe a positive correlation of activity of cluster.TEs with the expression of the genes they reside in. This was not observed generally for all gene TEs (data not shown). These findings could suggest that ‘cluster.TEs’ have a particular role in driving subtype-specific transcription profiles. Future work involving for example chromatin accessibility profiling and transcription start site sequencing (TSS-seq) would be valuable to further inform on this observation.

Tumor-specific TE-derived transcripts and peptides have been suggested as molecular tumor markers in various cancers [76]–[78]. Although retrotransposons in the human genome, including LTR and LINE1 elements, have undergone near-complete immobilization by recombination and mutation, they still retain in part translational competence [68]. We here describe a number of ATRT-subtype specific TEs with predicted coding capacity. Not only could these potentially serve as tumor-biomarkers, but might also be harnessed for tumor-directed immunotherapies as recently demonstrated in the context of renal cell carcinoma [79]. Immunogenicity of MYC-ATRTs has in addition previously been linked to sensing of double-stranded TE-derived RNAs [37]. We observe considerably higher levels of antisense transcription on LTR and LINE1 loci compared to human genes in primary ATRT samples (data not shown). Intriguingly, the pattern of bidirectional TE transcription appears to differ between ATRT subtype and TE family. MYC subtypes stood out with significantly lower bidirectionally transcribed LTR and LINE1 loci, however in particular for LTR elements, transcriptional activity of these selected loci appears to compensate in terms of overall TE-derived bidirectional transcripts compared to SHH and TYR subtypes. This might indicate that bidirectional TE expression is a targeted activity in MYC tumors. Of note, MYC expression itself has been demonstrated to be influenced by sense and antisense short RNAs transcribed from both strands either upstream or overlapping the first exon of the gene [80]. Moreover, MYC can regulate bidirectional promoters, such as *Pole3* [81]. Hence MYC may be driving specific bidirectional TE expression and it remains to be investigated if and which MYC-specific cellular pathways are involved.

To model ATRT tumorigenesis a number of cell line models have been generated, mimicking transcriptional and epigenomic features of ATRT-MYC or ATRT-SHH subtypes [26]. Our data confirm the heterogeneity of these models and marked differences concerning their transcriptomic profile compared to primary tumor samples. We here demonstrate that this also holds true for their genome-wide pattern of TE activity. Therefore, conclusions drawn from these cell lines regarding TE activity and TE-induced cellular effects might not be readily transferable to clinical situations. However, our data also suggest that general TE transcription characteristics observed in primary data are reflected in cell line data. For example, ATRT-MYC shows lower LINE1 transcription levels and less active LINE1 as well as LTR loci compared to ATRT-SHH samples. We furthermore show that expression of signature TE loci is sufficient to stratify tumor subtype identity in these models and therefore, as has been shown in one previous occasion [69], these models can to some extent be used to study the role of TEs in ATRT tumorigenesis in the future.

As we confined the study to ATRT samples with matching DNA-methylation and transcriptome data, a limitation of our study is the low sample size. Future prospective analyses of larger cohorts of ATRT subtype tumors are thus planned to confirm the here presented results.

Taken together, our study highlights molecular diversity of ATRT subtypes at the so far largely unexplored level of transposable element activity. It provides a comprehensive resource for future studies on TE implication in ATRT tumorigenesis, and calls for further exploration particularly in context of identification of novel ATRT biomarkers and optimization of immunotherapeutic approaches towards this cancer entity.

## Conclusions

Our data presents the first comprehensive transcriptional profiling of transposable elements (TEs) at locus-level resolution in primary atypical teratoid/rhabdoid tumors (ATRTs). The genome-wide TE activity profiles, define ATRT-MYC,-SHH and - TYR subtypes, reflecting their classification based on DNA methylation data. ATRT subtypes are further characterized by LTR and LINE1-specific signature loci, paving the way for detection of novel subtype specific treatment vulnerabilities. Specifically, ATRT-MYC tumors showed a distinct profile with specific bidirectional TE expression patterns, setting these tumors apart from SHH and TYR ATRTs. Finally, analyses of the TE transcriptome in ATRT cell line models revealed that they partly reflect the TE patterns observed in primary ATRTs, setting a basis for future functional studies. As TEs are suspected to drive ATRT immunogenicity, our data sets the foundation for novel therapeutic interventions, such as immunotherapy.

## List of abbreviations

ATRTs: Atypical teratoid rhabdoid tumors
ATRT MYC: Atypical teratoid rhabdoid tumor, MYC subtype
ATRT SHH: Atypical teratoid rhabdoid tumor, Sonic Hedgehog subtype
ATRT TYR: Atypical teratoid rhabdoid tumor, tyrosinase subtype
DE: Differentially expressed
DEG: Differentially expressed genes
LINE1: Long interspersed nuclear elements 1
LTR: Long terminal repeats
TE: Transposable element

## Declarations

Ethics approval and consent to participate

Not applicable

## Consent for publication

Not applicable

## Availability of data and material

The datasets analyzed during the current study are available on the Gene Expression Omnibus (GEO) repository, https://www.ncbi.nlm.nih.gov/geo/ under the accession numbers: GSE231287, SRP457328 & SRP322114.

## Authors’ contributions

MVH and SG analyzed data and wrote the manuscript. MA, SML and JN analyzed and interpreted data and contributed to figures, GT and DJM generated cell line gene expression data and analyzed data, UCL and JEN supervised the study, interpreted data and wrote the manuscript.

## Acknowledgements

We thank the laboratory teams of JEN and UCL for excellent technical support and helpful discussions, especially Tasja Lempertz, Paula Rieck, Antonia Gocke, Yannis Schumann and Matthias Dottermusch.

---

**Suppl. Fig. 1: Signature gene expression analysis in primary ATRT samples**

Pairwise differential gene expression (DEG) analysis between all three ATRT subtypes (MYC, SHH and TYR). Cutoffs for DEGs set to log2FC >|2|, padj < 1×10E-06. a) Unsupervised hierarchical clustering of all differentially expressed genes combined. b) Unsupervised hierarchical clustering of 42 ATRT signature genes being de-regulated in specific ATRT subtypes.

**Suppl. Fig. 2: Sample clustering based on gene and TE transcription profile subsets**

a) Representation of TE bins with respect to gene transcriptional start sites. b) Principal component analysis of ATRT samples based on gene and TE transcription, respectively. PCA based on vst-transformed counts, plotted 500 most variable genes/TEs.

**Suppl. Fig. 3: Differences in TE transcription at subfamily level (extended from Fig. 2)**

Relative normalized TE counts were summarized at TE subfamily level, excluding elements with less than 5 mapped reads on average in at least one ATRT subgroup (expression score). Occurrence score is defined as the number of expressed loci per subfamily [in %]. Shown are the three LTR families ERV1, ERVL and ERVL-MaLR.

