## Supplementary figures and images for "Expression of LTR and LINE1 transposable elements defines atypical teratoid/rhabdoid tumor subtypes"

### Supplemental figure 1

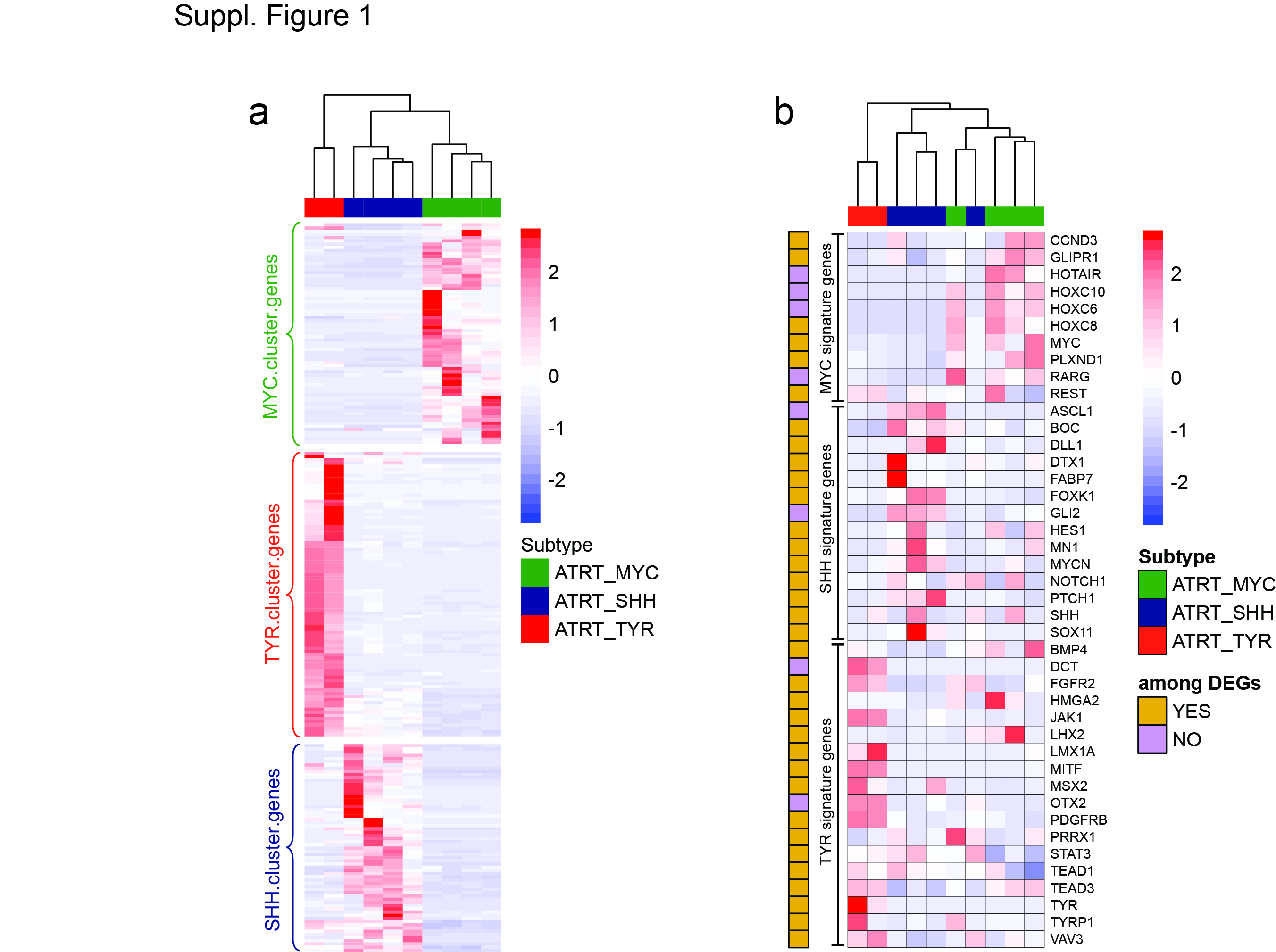

### Supplemental figure 2

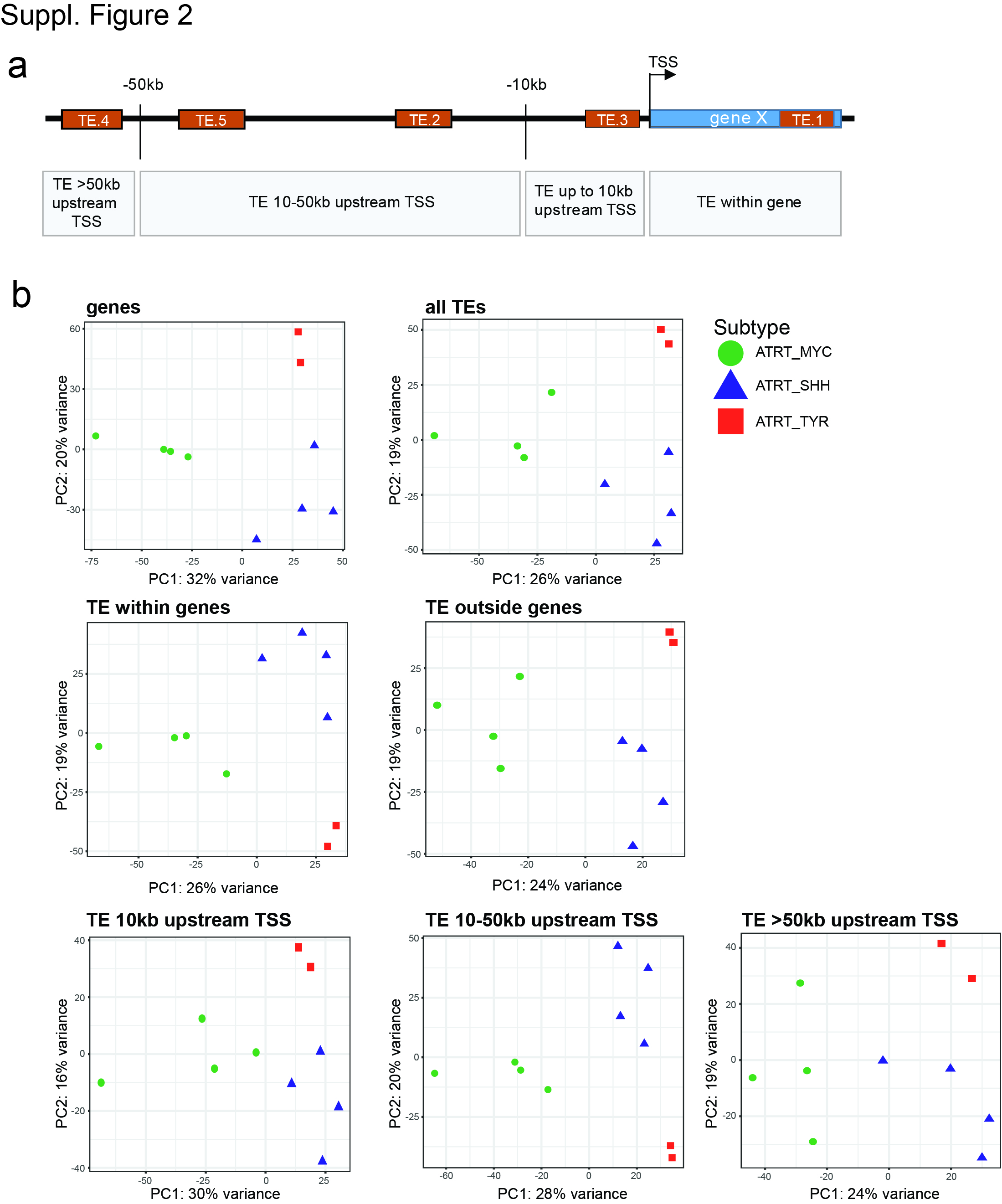

### Supplemental figure 3

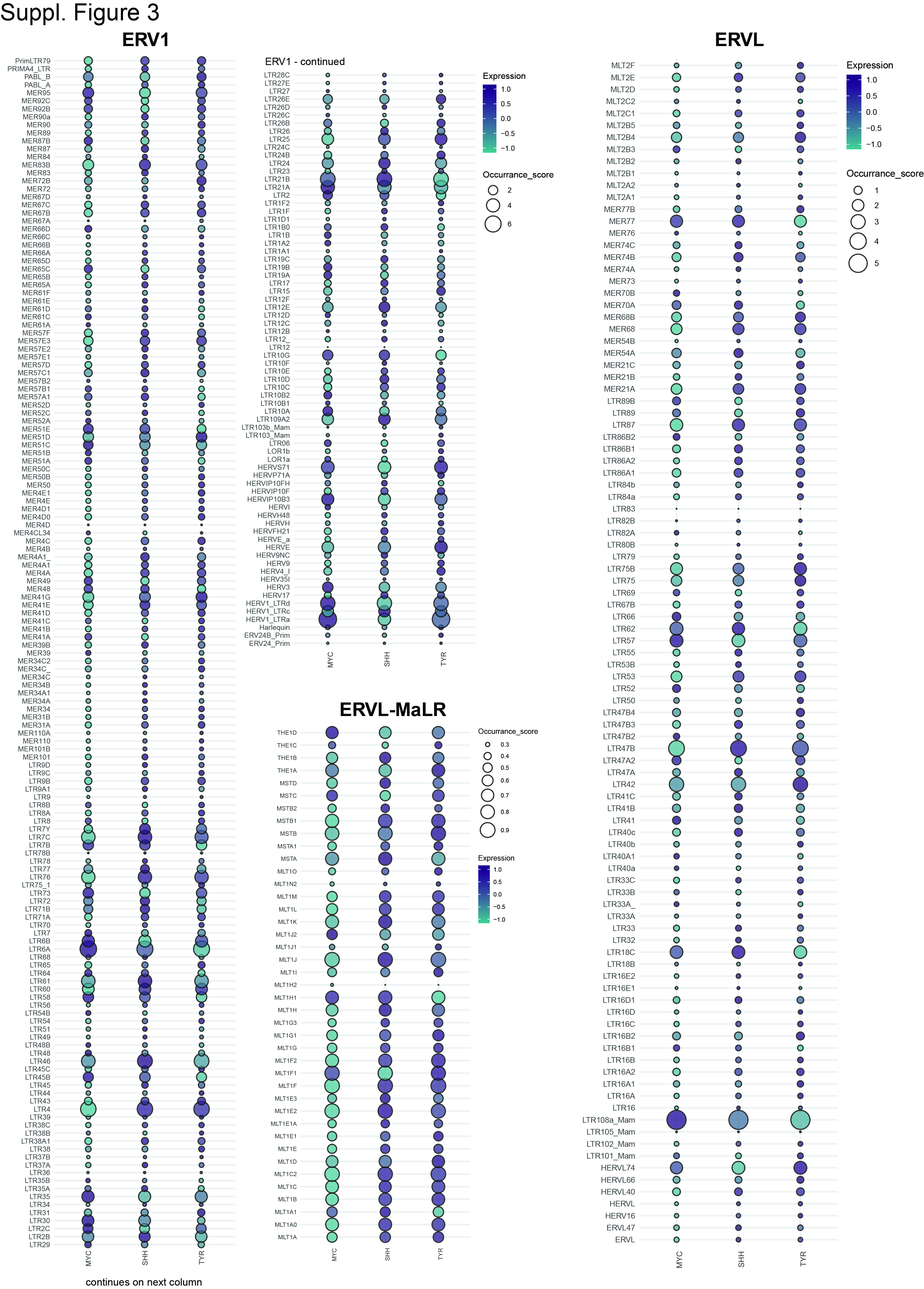
